# Modulation of sleep by trafficking of lipids through the Drosophila blood brain barrier

**DOI:** 10.1101/2022.02.17.480875

**Authors:** Gregory Artiushin, Fu Li, Amita Sehgal

## Abstract

Endocytosis through Drosophila glia is a significant determinant of sleep amount and occurs preferentially during sleep in glia of the blood brain barrier (BBB). To identify metabolites whose trafficking is mediated by sleep-dependent endocytosis, we conducted metabolomic analysis of flies that have increased sleep due to a block in glial endocytosis. We report that acylcarnitines, molecules that conjugate with long chain fatty acids to promote their transport, accumulate in heads of these animals. In parallel, to identify transporters and receptors whose loss contributes to the sleep phenotype caused by blocked endocytosis, we screened genes enriched in barrier glia for effects on sleep. We find that knockdown of lipid transporters *LRP1*&2 as well as carnitine transporters *ORCT1*&2 increases sleep. In support of the idea that the block in endocytosis affects trafficking through specific transporters, knockdown of LRP or *ORCT* transporters also increases acylcarnitines in heads. We propose that lipid species, such as acylcarnitines, are trafficked through the BBB via sleep-dependent endocytosis, and their accumulation in the brain increases the need for sleep.

## Introduction

Although the regulation of sleep is normally studied on a behavioral and circuit level, there is increasing evidence for a role of basic cellular physiology. For instance, we found that disruption of endocytic and trafficking pathways in glia increases sleep in Drosophila (Artiushin, Zhang et al. 2018). Glia of the Drosophila blood/hemolymph brain barrier (BBB) emerged as a new cellular locus of sleep regulation in this study, such that genetically manipulating endocytosis in these cells alone was sufficient to increase sleep. As the increased sleep appeared to reflect higher sleep need, we asked if sleep is typically required for endocytosis through the BBB and found that this was indeed the case. However, the nature of the molecules trafficked through the BBB in a sleep-dependent manner was not known.

Barriers that separate the solutes of blood/hemolymph of the periphery from the interstitial fluid of the central nervous system display a rich profile of transporters, receptors, and trafficking proteins, which often reflect their unique functions. The *Drosophila* barrier glia populations share many conserved features with vertebrate barriers which employ endothelial and astrocytic populations (DeSalvo, Hindle et al. 2014, Weiler, Volkenhoff et al. 2017). For instance, both are capable of moving lipids and carbohydrates, ions, amino acids and xenobiotics (Weiler, Volkenhoff et al. 2017). Furthermore, the fly barrier populations may serve specialized roles in metabolism, as not only the conduit of energy sources from the periphery, but also by containing the enzymatic machinery necessary for processing energy sources (Volkenhoff, Weiler et al. 2015), and secreting signals in reference to nutritional state (Chell and Brand 2010, Speder and Brand 2014).

Given that much of the traffic through the barrier involves energetic substrates, we conducted metabolomic profiling to identify candidate metabolites whose trafficking may be inhibited by an endocytosis block in glia. To complement this approach, we asked if specific glial proteins mediate this trafficking. As genetic manipulations to block endocytosis can directly impact endocytosis-dependent carrier traffic as well as indirectly affect levels of membrane-associated transporters/receptors by altered recycling, we performed a knockdown screen to broadly search for barrier genes involved in sleep regulation. We report here that specific lipid and carnitine transporters act in barrier glia to affect sleep, and that disrupting expression of these transporters or of endocytosis leads to an accumulation of acylcarnitines in the head.

## Results

### Expression of *Shi* in glia induces acylcarnitine accumulation in fly heads

Our previous work showed that expression of *shibire (shi*), a dominant negative dynamin that blocks endocytosis, in all or BBB glia increases sleep. To attain an unbiased, global assessment of metabolites that may be relevant to the increased sleep seen in *Repo>20xShi.ts1* flies (hereafter referred to as *Repo>Shi^1^*), we conducted LC-MS analysis. Heads of male and female *Repo-Gal4 > Shi^1^* flies as well as Gal4 and UAS controls were collected on dry ice with sieves and immediately frozen at −80 ℃. Each sample contained 200 fly heads (equal male and female), with a total of 5 samples per genetic condition.

As an initial analysis, raw signal was scaled per each metabolite in reference to other samples within the dataset, and comparisons were made between Repo>Shi flies and each control, as well as controls to each other by Welch’s t-test (**Supplementary Table 1**).

Metabolites of interest were those for which signal from the experimental samples was significantly different, in the same direction, when compared to both controls, while controls compared to each were not significant. Of secondary interest were metabolites where a difference was seen in controls, but was proportionally smaller than consistent differences of each control to the experimental samples.

In surveying this dataset, the outstanding functional category, which contained multiple metabolites whose signal was consistently different in experimental animals versus controls, were the acyl-carnitines (**Table 1**). Furthermore, the fold changes for given metabolites in this group, which consists of fatty acids conjugated to carnitine, were the highest overall. Carnitinylation occurs on fatty acids of various chain lengths, but only a subset of chain lengths in this dataset had sufficient signal, therefore we statistically compared *Repo>Shi^1^* flies to both parental controls for metabolites that had signal in at least three of five biological replicates for each genotype. Expression of *Shi^1^* in glia increased abundance in fly heads of the following acylcarnitine species: C2, C16, C16*1, C17, C18:1, C18:2* (**Figure 1A**). The only metabolite of this group with less signal in experimental animals was the longer-chain, C24* (**Figure 1B**). Carnitine and deoxycarnitine were not significantly altered as compared to both controls.

**Table 1.** Fatty acid acylcarnitine accumulation in *Repo*>*Shi^1^* fly heads. All samples from *Repo-GAL4>UAS-Shi^1^*, and both parental controls. Welch’s t-test was performed on scaled signal for each metabolite, comparing the conditions shown. Green highlighting marks a significant difference (p≤0.05) between the groups, where metabolite ratio is < 1.00, while light green is not significant, but close to the threshold (0.05<p<0.10). Red highlighting marks a significant difference (p≤0.05) between groups where metabolite ratio is ≥ 1.00, and light red is not significant, but close to the threshold (0.05<p<0.10).

| Sub pathway | Biochemical name | $\frac{Gal4 > UAS}{Gal4Ctrl}$ | $\frac{Gal4 > UAS}{UASCtrl}$ | $\frac{UASCtrl}{Gal4Ctrl}$ |
| --- | --- | --- | --- | --- |
| Fatty Acid Metabolism (Acylcarnitine) | acetylcarnitine (C2) | 4.25 | 2.68 | 1.59 |
|  | myristoylcarnitine (C14) | 28.81 | 26.39 | 1.09 |
|  | palmitoylcarnitine (C16) | 4.52 | 1.74 | 2.60 |
|  | palmitoleoylcarnitine (C16:1) | 26.50 | 10.35 | 2.56 |
|  | stearoylcarnitine (C18) | 1.43 | 1.06 | 1.34 |
|  | linoleoylcarnitine (C18:2) | 6.77 | 3.14 | 2.15 |
|  | oleoylcarnitine (C18:1) | 4.00 | 1.92 | 2.08 |
|  | arachidoylcarnitine (C20) | 1.28 | 1.17 | 1.09 |
|  | behenoylcarnitine (C22) | 1.31 | 1.09 | 1.20 |
|  | eicosenoylcarnitine (C20:1) | 2.28 | 1.24 | 1.84 |
|  | lignoceroylcarnitine (C24) | 0.72 | 0.83 | 0.87 |
|  | margaroylcarnitine (C17) | 4.05 | 1.47 | 2.76 |
|  | nervonoylcarnitine (C24:1) | 2.36 | 1.26 | 1.87 |
|  | cerotoylcartinine (C26) | 0.85 | 0.97 | 0.87 |
|  | ximenoylcarnitine (C26:1) | 0.75 | 0.88 | 0.85 |

**Figure 1:**
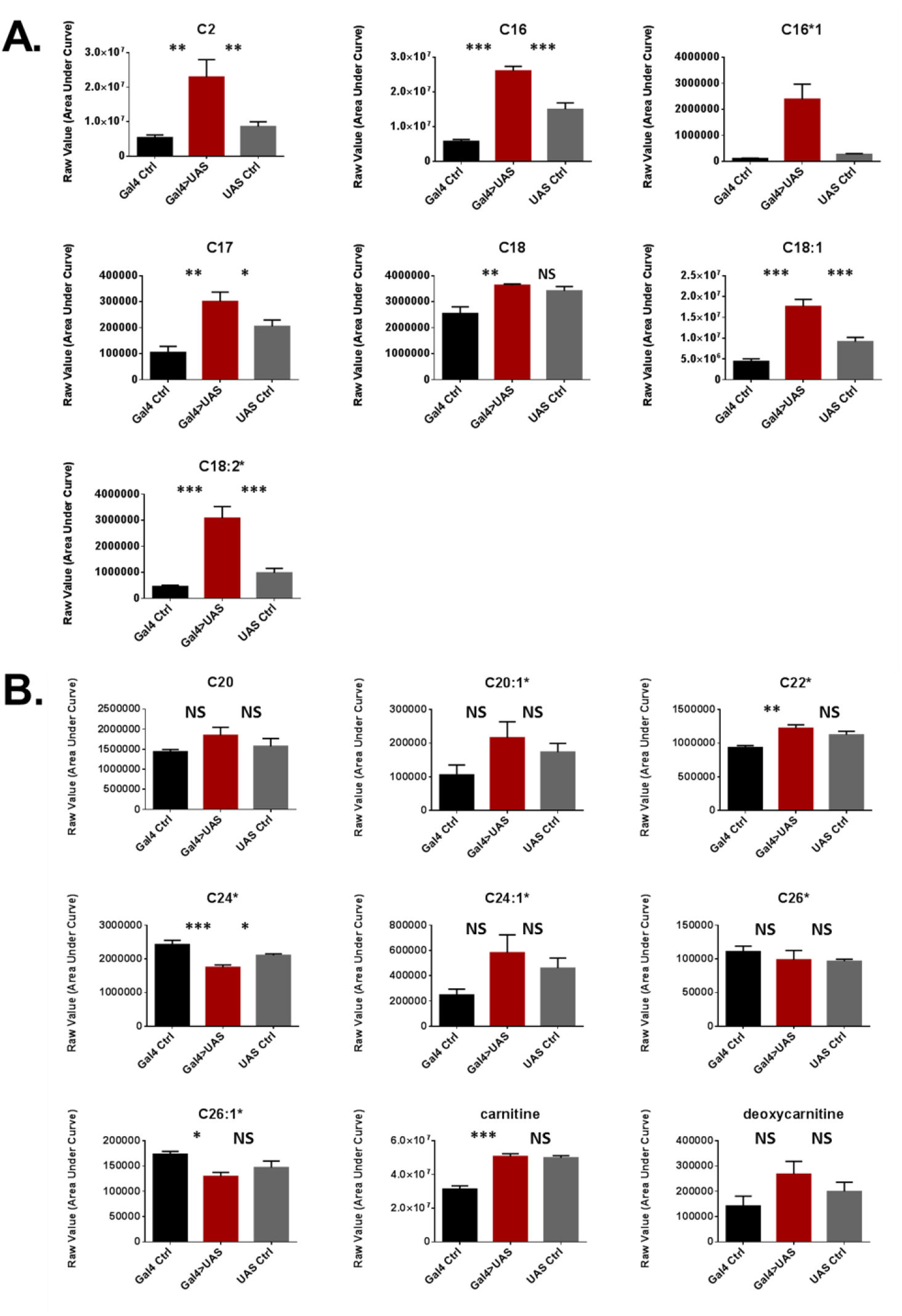
Acylcarnitine levels are increased in *Repo>Shi^1^* fly heads. (**A**) Short and medium chain length or (**B**) long chain length acylcarnitines from Repo-G4 >UAS-Shi^1^ fly heads and parental controls. The raw signal from LC/MS is plotted, n=3 – 5 samples, of 200 fly heads each. One-way ANOVA, with Holk-Sidak post-hoc comparisons. *p < 0.05, **p < 0.01, ***p < 0.001. Error bars represent standard error of the mean (SEM).

### Identification of barrier glia genes that affect sleep

In addition to identifying metabolites that accumulate as a result of blocked glial endocytosis, we sought to identify glial molecules whose function might be impacted by the block in endocytosis and thereby contribute to the effect on sleep. As noted above, glia of the BBB are the most relevant glial subtype for sleep-dependent endocytosis. We identified genes enriched in barrier glia by referring to transcriptional profiling that compared expression in the two glial populations that comprise the Drosophila BBB— subperineurial and perineurial glia (SPG + PG)— to all neurons, and all glia (DeSalvo, Hindle et al. 2014). Preference was given to previously studied genes, particularly transporters, receptors and those involved in trafficking, although many genes among the top-50 highly expressed in the barrier glia populations were also tested for effects on sleep. Of the genes enriched in barrier glia, we focused on those that showed low variability in expression from sample to sample. UAS-RNAi constructs for candidate genes were expressed with *Repo/RepoGS* Gal4 drivers and sleep in these lines was compared with that in Gal4 and UAS alone controls. Knockdown of most genes did not produce a significant phenotype, but sleep was increased with knockdown of some transporter genes *CG3036*, *CG6126*, *mnd, VMAT, CG6836, Rh50, CG4462* (**Figure 2- figure supplement 1A**), cytoskeleton/trafficking factors *CG8036*, *Vha16, nuf*, (**Figure 2-figure supplement 1C**) as well as *lsd-2*, *acon* and *MtnA* (**Figure 2- figure supplement 1D**). Meanwhile, knockdown of the transporter gene *CG16700* (Supp Figure 2A), cytochrome P450 gene *Cyp6a20* (**Figure 2- figure supplement 1D**) and trafficking factor *Cln7* decreased total sleep (**Figure 2- figure supplement 1C**).

**Figure 2:**
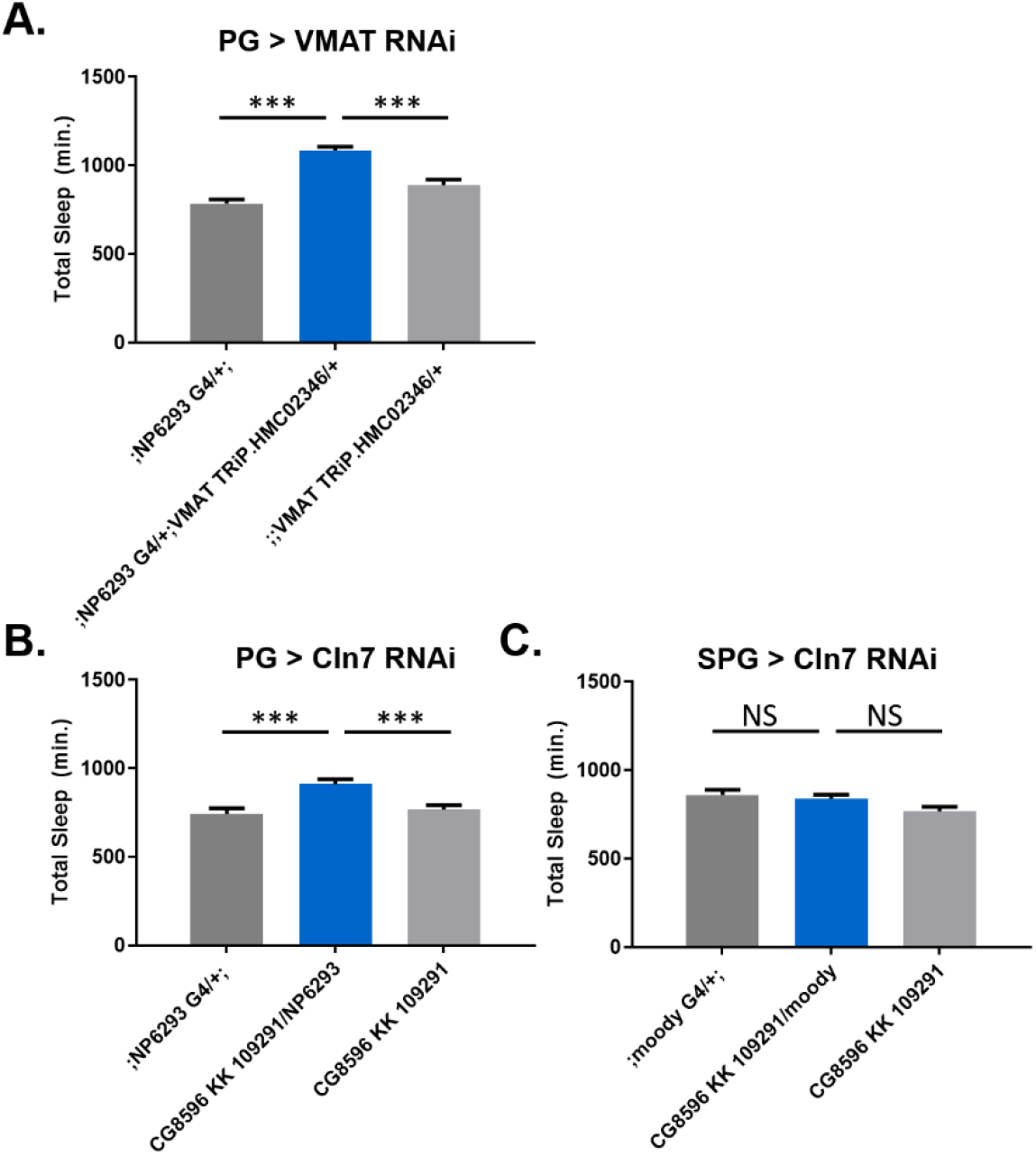
Knockdown of specific barrier-enriched genes with barrier glial drivers. Total sleep in female flies with knockdown of (**A**) VMAT, UAS-HMC02346 driven by (PG) NP6293-Gal4. n=15-16 per genotype. (**B and C**) Cln7 (CG8896), UAS-109291 KK driven by (PG) NP6293-Gal4 or (SPG) moody-Gal4. n = 15-16 per genotype. One-way ANOVA, with Holk-Sidak post-hoc comparisons. *p < 0.05, **p < 0.01, ***p < 0.001. Error bars represent standard error of the mean (SEM).

Since the candidate genes were selected based on enrichment within the barrier glia, we chose to examine and secondarily validate promising phenotypes through knockdown with more limited, barrier glia drivers. Thus, we screened a sub-set of the genes suggested by results of the pan-glial screen (**Figure 2-figure supplement 1)** with drivers that target PG or SPG glia. Knockdown of MtnA (105011 KK), CG6386 (108502 KK), CG4462 (105566 KK), or lsd-2 (102269 KK) did not significantly alter sleep when expressed in either of the barrier glial populations alone (data not shown). Reduction of *cyp6a20* in the PG population inconsistently reproduced the pan-glial sleep loss phenotype, so this was not pursued further (data not shown). The *VMAT* gene produces two isoforms, one of which is thought to be specific to glia (Romero-Calderon, Uhlenbrock et al. 2008). Knockdown of *VMAT* (TRiP HMC02346) in the perineurial glia increased total sleep (**Figure 2A**), but in the subperineurial glia it had no effect (data not shown), which is consistent with protein expression, as the *VMAT-B* antibody specifically marks the perineurial glia (DeSalvo et al., 2014). Pan-glial knockdown of neuronal ceroid lipofuscinosis 7 (*Cln7*), which is expressed in the perineurial glia (Mohammed et al., 2017), decreased total sleep (**Figure 2C**). However, expression of *Cln7* RNAi in the PG increased total sleep time, with no significant change produced by expression in the SPG (**Figure 2B, C**).

### Simultaneous knockdown of *Lrp1* and *Megalin* in barrier glia increases sleep

Although the candidate screen identified barrier genes whose knockdown increases sleep, as does blocking endocytosis in barrier glia, these genes were not obviously linked to the metabolite profile seen with a block in glial endocytosis. The results of metabolomic screening showed changes in lipid, and particularly carnitine-lipid, trafficking. Therefore, we reassessed our screen candidates to consider transporters and receptors which may function in these pathways and could have been missed due to redundancy/lethality. *Lrp1* and *Megalin* (*Lrp2*) are two LDL receptor-related protein members involved in the transport of lipid carrier proteins at the fly barrier (Brankatschk, Dunst et al. 2014). Expression is likewise found in mammals at the endothelial barrier (Herz 2003).

Knocking down *Lrp1* and *Megalin* (*Lrp2*) individually in the pan-glial screen did not significantly alter total sleep time (**Figure 2- figure supplement 1A**). However, both *Lrp1* and *Megalin* (Lrp2) as well as *Orct* and *Orct2* have been considered to be complementary (Eraly and Nigam 2002, Brankatschk, Dunst et al. 2014), therefore it is possible that inhibition of a single gene is insufficient to appreciably affect transport. Simultaneously knocking down *Lrp1* and *Megalin* in all glia with Repo-Gal4 driver increased total sleep time (**Figure 3A).**To target *Lrp1* and *Megalin* in barrier glia, and also to restrict the knockdown to the adult stage and thereby avoid developmental confounds, we used drivers specific to these glia and coupled them with the temperature-sensitive tubulin-Gal80 (tub-Gal80ts) system that suppresses Gal4 expression at 18 degrees but allows it at 31 degrees. When both *Lrp* genes were knocked down with barrier glia drivers, a significant increase in total sleep was seen with the PG driver (NP6923), but not with either of the SPG drivers (Moody, Rab9) (**Figure 3B, C, D, E**). Knockdown of *Lrp1* and *Megalin* with the 9-137 driver, which expresses in both PG and SPG, also increased total sleep significantly (**Figure 3C and D**). Intriguingly, although the Repo driver did not yield a phenotype with *Lrp1* alone, each of two *Lrp1* RNAi constructs (GD 8397, GD 13913) expressed by the driver 9-137-Gal4 increased total sleep (**Figure 3- figure supplement 1**).

**Figure 3.**
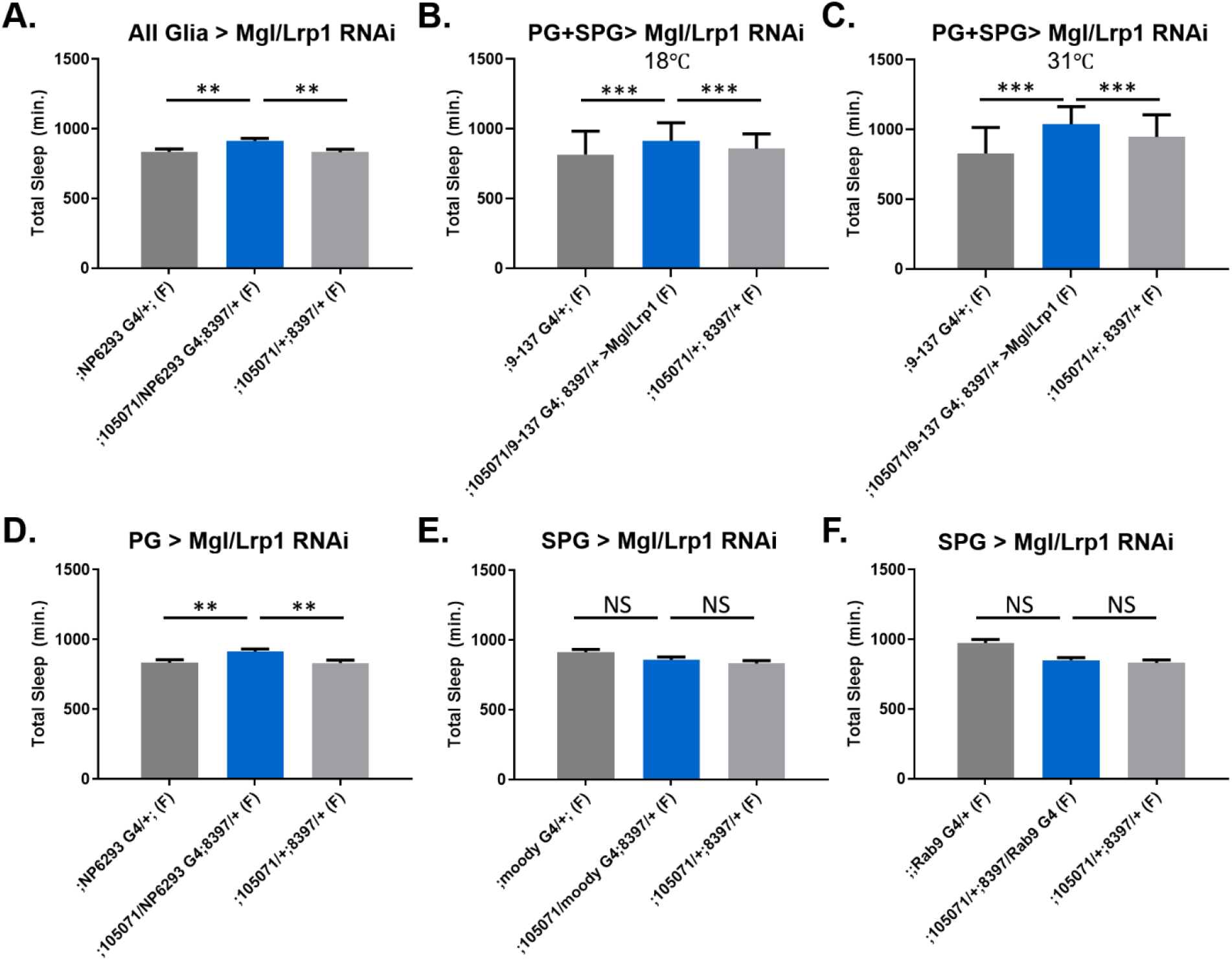
Sleep time changes with knockdown of Lrp genes in all glia or barrier glia. Total sleep in female flies with knockdown of (**A**) *Lrp1* (8397 GD) and *Megalin* (105071 KK) RNAi driven by Repo-GAL4. n = 11-16 per genotype; (**B**) *Lrp1* (8397 GD) and *Megalin* (105071 KK) RNAi driven by 9-137-GAL4 (PG and SPG) with TubGal80ts at the permissive temperature of 18 ℃, n=30- 32 per genotype; (**C**) *Lrp1* (8397 GD) and *Megalin* (105071 KK) RNAi driven by 9-137-GAL4 (SPG and PG) with TubGal80ts at the restrictive temperature of 31 ℃, n=30- 32 per genotype.; (**D, E, F**) *Lrp1* (8397 GD) and *Megalin* (105071 KK) RNAi expressed by: (D) NP6293- Gal4 (**PG**), n = 13-16 per genotype; (**E**) *moody*-GAL4 (SPG), n=16 per genotype; (**F)** Rab9-GAL4 (SPG), n=13-16 per genotype. One-way ANOVA, with Holk-Sidak post-hoc comparisons. *p < 0.05, **p < 0.01. Error bars represent standard error of the mean (SEM).

### Knockdown of *Orct* and *Orct2* in barrier glia increases sleep

The organic cation (*Orct*) transporters are multi-substrate transporters whose substrates include carnitine (Lahjouji, Mitchell et al. 2001), and perhaps also carnitylated molecules, based on *in vitro* evidence for *Orct2* (Kou, Yao et al. 2017). As with the double knock down of *Lrp1* and *Megalin*, knockdown of both *Orct* genes increased total sleep (**Figure 4A**). In fact, for the *Orct* genes, knockdown in either PG or SPG replicated the increased total sleep phenotype, although this was only true with one line for the SPG (**Figure 4B**). It is worth noting that the phenotypes with the barrier drivers are considerably more moderate than with knockdown in all glia.

**Figure 4.**
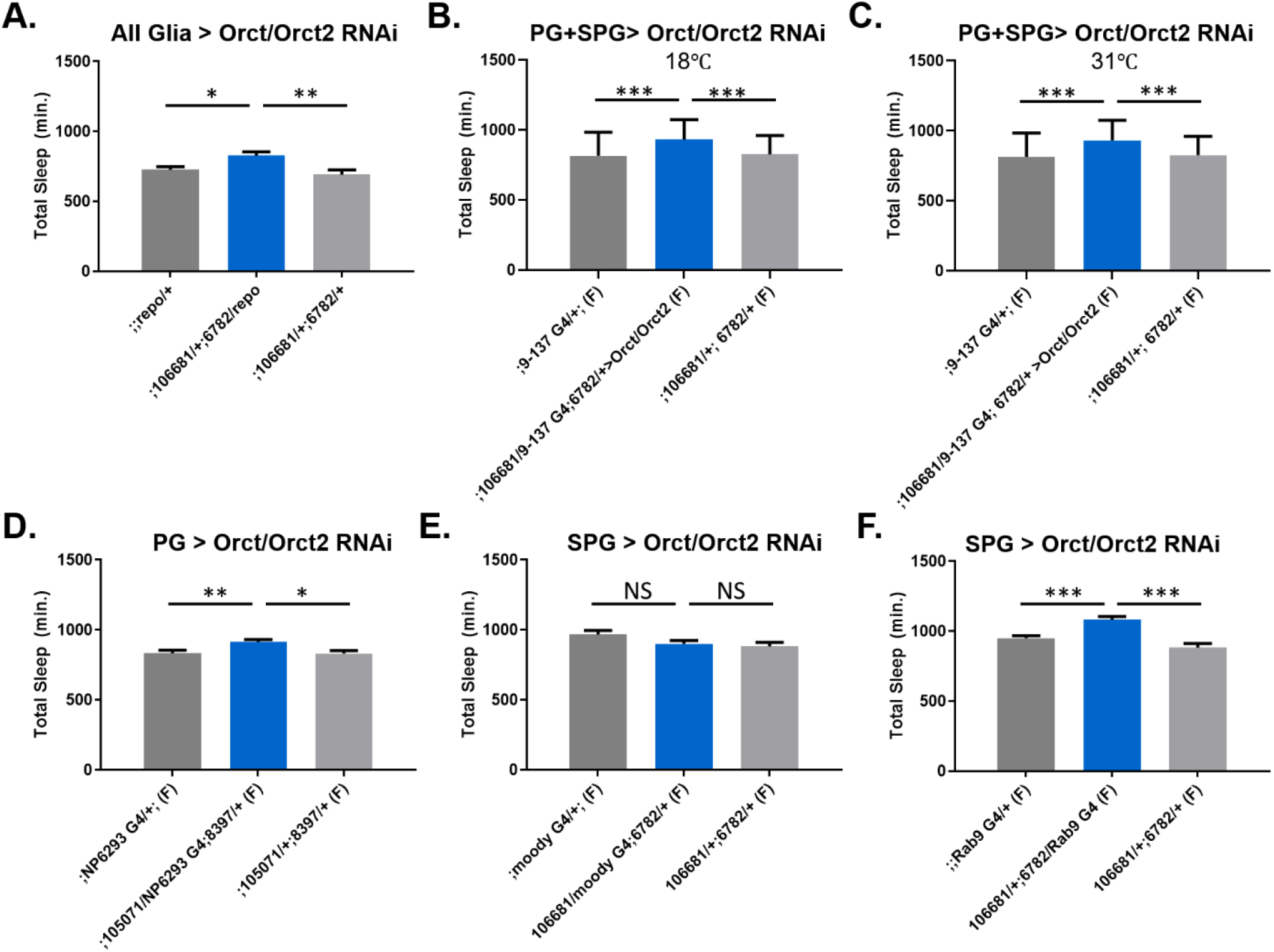
Sleep time changes with knockdown of *Orct* genes in all glia or barrier glia. RNAi constructs of *Orct1* and *Orct2* were expressed as follows: (**A)** *Orct* (6782 GD) and *Orct2* (106681 KK) driven by Repo-GAL4; (**B and C)** *Orct* (6782 GD) and *Orct2* (106681 KK) driven by (PG+SPG) 9-137-Gal4 at 18 ℃(permissive), n=30- 32 per genotype or at 31℃(restrictive), n=30 - 32 per genotype; (**D-F**) *Orct* (6782 GD) and *Orct2* (106681 KK) driven by NP6293-GAL4, n = 13-16 per genotype or by (SPG) *moody*-GAL4, n=15-16 per genotype or by (SPG) Rab9- GAL4, n=15-16 per genotype. One-way ANOVA, with Holk-Sidak post-hoc comparisons. *p < 0.05, **p < 0.01, ***p < 0.001. Error bars represent standard error of the mean (SEM).

### Knockdown of Lrp and *Orct* genes in glia leads to accumulation of acylcarnitines

Given that knockdown of the *Lrp* and *Orct* genes in glia parallels effects of *Shi* expression in terms of increasing total sleep, we asked if it had the same effect on metabolite accumulation in fly heads. Based on the metabolomic results of *Repo>Shi^1^*, we chose to specifically assay acylcarnitines through LC-MS analysis. Metabolomic profiling requires considerable tissue, as so, as in the case of the *Shi^1^* experiment, we collected heads from flies in which either *Lrp1* and *Megalin* or *Orct 1* and *Orct2* were knocked down with *Repo*-Gal4.

Knockdown of *Lrp1* and *Megalin* in all glia increased abundance in fly heads of the following acylcarnitine species: C3, C4 butyryl, C4-OH isobutyryl, C4 isobutyryl, C16, C18, C20 (**Figure 5**). Longer-chain acylcarnitines, e.g. those over C22, were largely undetected in experimental samples. Knockdown of *Orct* and *Orct2* similarly enriched acylcarnitines in fly heads. In particular, acylcarnitine species C16, C16:1 and C20 were increased significantly compared to their controls (**Figure 6**). Similar in the case of the *Lrp* samples, longer-chain acylcarnitines over 22 were not detected well.

**Figure 5.**
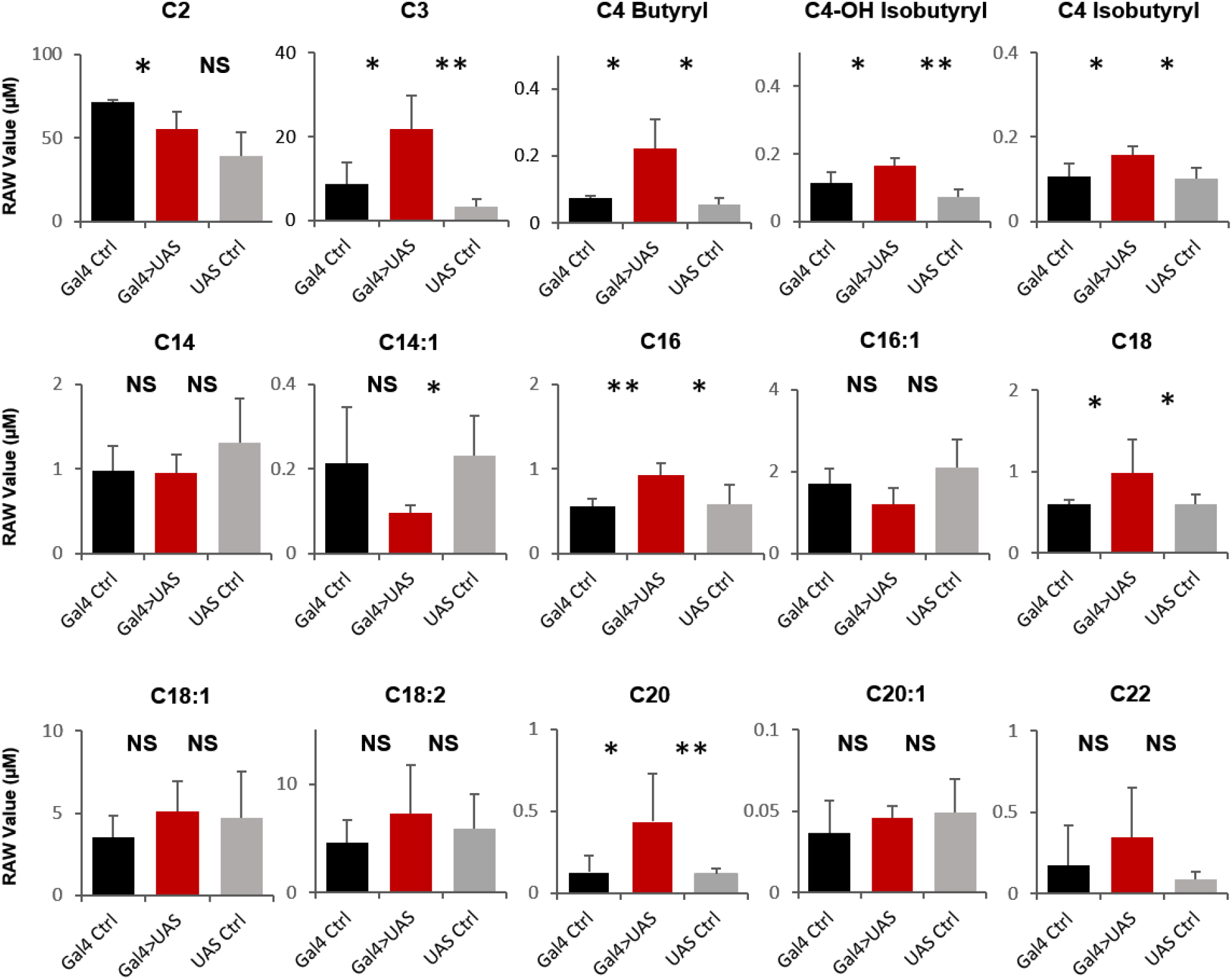
Acylcarnitine levels are increased in Repo>*Lrp1*+ Mgl fly heads. Short and medium chain length acylcarnitines from Repo Gal4 >*Lrp1*+Mgl fly heads and parental controls. The raw signal from LC/MS is plotted, n=3 samples, of 300 fly heads each. Student's t-test with comparison. *p < 0.05, **p < 0.01. Error bars represent standard error of the mean (SEM).

**Figure 6.**
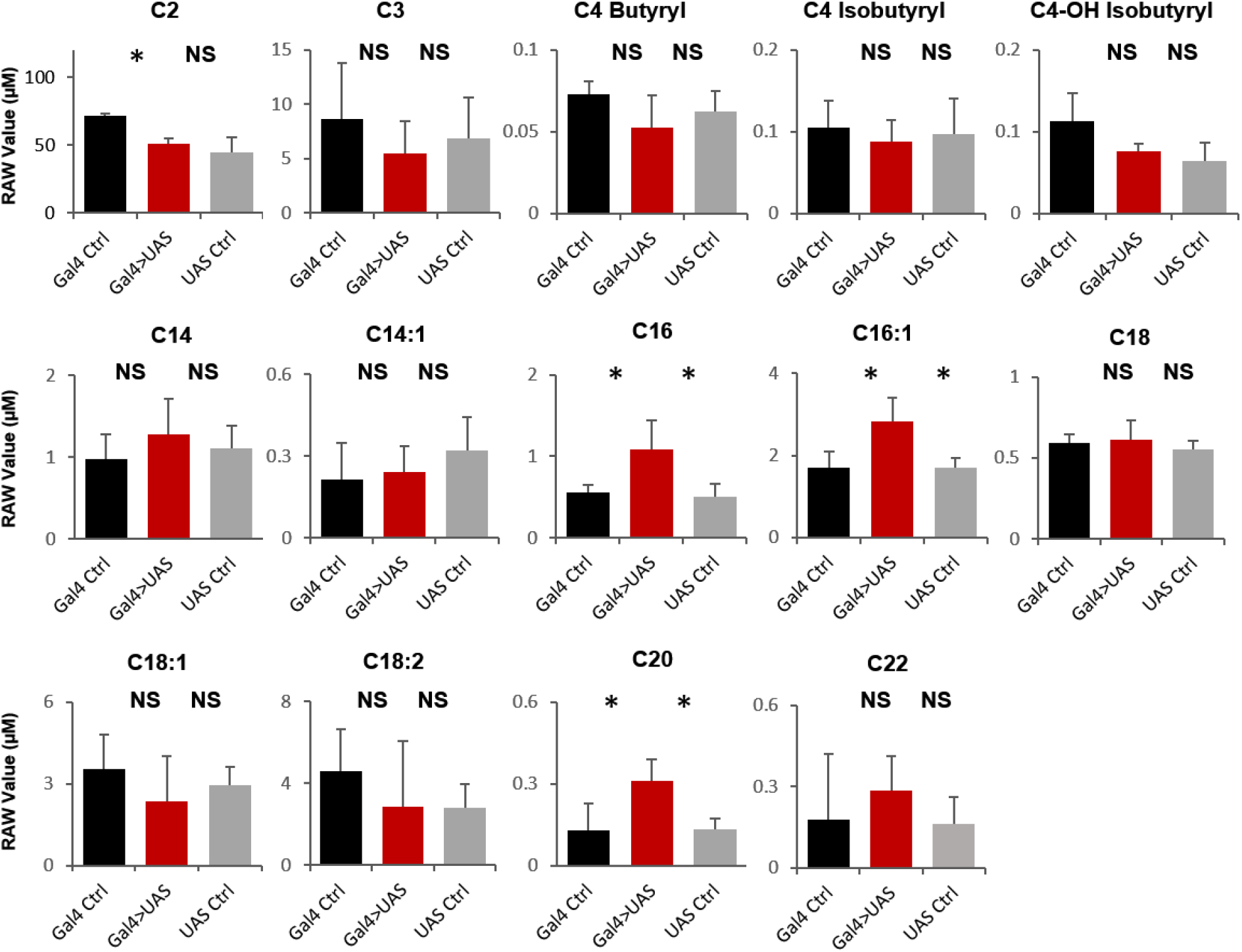
Acylcarnitine levels are increased in Repo> *Orct*+*Orct2* fly heads. Short and medium chain length acylcarnitines from *Repo-Gal4 >Orct+Orct2* fly heads and parental controls. The raw signal from LC/MS is plotted, n=3 samples, of 300 fly heads each. Student's t-test with comparisons. *p < 0.05, **p < 0.01. Error bars represent standard error of the mean (SEM).

## Discussion

Our previous work suggested that endocytosis through the BBB is a function of sleep, but the nature of the molecules trafficked remained unknown. Using a dual-pronged approach of a targeted genetic screen and unbiased metabolomic analysis, we report here that the passage of lipids through the BBB is important for sleep. Blocking such transport increases sleep in conjunction with an accumulation of acylcarnitines.

Through a pan-glial RNAi knockdown screen of candidate genes expressed in the fly barrier, we identified molecules that affect daily sleep amount. In follow-up experiments, we targeted knockdown to each barrier layer separately, which is subject to the concern that behavioral phenotypes requiring simultaneous knockdown in both barrier populations would be missed. Nevertheless, we consider this scenario to be less probable, as permeability through the populations is quite different, with smaller solutes likely passing through the PG but not the tight barrier of the SPG, lipophilic solutes or xenobiotics potentially passing through each uninhibited, and larger solute requiring endocytic mechanisms that likely have to work in each population in tandem. Therefore, in most cases of knockdown, transport would either be inhibited by the one barrier population essential for it, or would be interrupted by either population. This was indeed the case for the lipid and carnitine transporters we focused on, where knockdown in a specific layer or either layer of the BBB was sufficient for a phenotype. While there was redundancy at the level of the transporters, the two barrier layers did not compensate for each other.

An additional consideration of preliminarily screening with pan-glial drivers is that if knockdown in multiple glial subtypes has opposing effects on sleep, we may have obscured a role for the barrier cells. Again, we attempted to minimize this risk by primarily selecting genes whose expression is both highly abundant and specifically enriched in the barrier populations, as opposed to the set of all glial cells in the transcriptome dataset (DeSalvo et al., 2014). The assumption is that multifold expression in the barrier populations is indicative of prevailing importance in these cells, although this is a caveat.

The vesicular monoamine transporter (VMAT) has previously been identified as a target of reserpine, which promotes sleep in the fly (Nall et al., 2014). VMAT mutants exhibit higher baseline sleep, and also lose less sleep than controls when subject to mechanical sleep deprivation. VMAT can traffic multiple monoamines such as dopamine, serotonin, histamine and octopamine, but no single neuronal population or neurotransmitter system was implicated as responsible for the VMAT sleep phenotype (Nall et al., 2014).

In flies, VMAT exists as two isoforms, VMAT-A, which is expressed in monoaminergic neurons, and VMAT-B, which appears to be specific to perineurial glia (DeSalvo et al., 2014), as it is also found in fenestrated glia in the visual system (Romero-Calderon, Uhlenbrock et al. 2008), which are a specialized form of perineurial glia (Kremer, Jung et al. 2017). It is unknown whether VMAT-B would functional similarly in glia as VMAT-A does in neurons. VMAT-B contains an additional cytoplasmic domain, which has been suggested to promote retention in the plasma membrane as opposed to trafficking to vesicles (Greer, Grygoruk et al. 2005). *VMAT* knockdown increased sleep, as did disrupting endocytic trafficking at the barrier (Artiushin, Zhang et al. 2018). Whole-brain levels of monoamines were not altered in flies that expressed *Shi^1^* in glia, nevertheless it is possible that this gross analysis would not be sensitive to local changes at the barrier. In the visual system glia, VMAT-B may be necessary for uptake of histamine (Romero-Calderon, Uhlenbrock et al. 2008). Interestingly, histamine is known to alter permeability of the blood-brain barrier in mammals (Lu, Diehl et al. 2010).

*Cln7* is a major facilitator superfamily transporter implicated in neuronal ceroid lipofuscinoses, and hence considered to impact lysosomal/autophagal function (Siintola, Topcu et al. 2007). Neither the function, nor what this transporter traffics, are known, but it is thought to be vesicular as well, and is expressed in the perineurial glia in flies (Mohammed, O'Hare et al. 2017). Knockdown of *Cln7* in all glia affected sleep in the opposite direction from knockdown only in the PG. One potential explanation would be that *Cln7* acts on sleep in opposing ways in different glial populations, although protein expression data suggest that *Cln7* is quite limited in the brain, and it is not clear whether it is in other glial populations.

Our metabolomic data indicated that acylcarnitines are elevated in the heads of the long-sleeping Repo>Shi^1^ flies. Acylcarnitines are transported to mitochondria for fatty acid oxidation, but are also secreted as they are found in plasma in mammals (Schooneman, Vaz et al. 2013). Given that circulating acylcarnitines can be taken up by cells, we investigated *Lrp1*/*Megalin* and *Orct*/*Orct2* as candidate transporters for this uptake. Lrp1 and Megalin are lipoprotein carrier receptors known to function in the fly barrier (Brankatschk, Dunst et al. 2014) and although knockdown of each one separately in all glia did not affect sleep, reducing expression of both in all glia or barrier glia increased sleep. Likewise, knockdown of *Orct* and *Orct2*, which are homologs of the human carnitine transporters (Eraly and Nigam 2002) and transport carnitine as well as acylcarnitines (Pochini, Oppedisano et al. 2004), increases sleep. LC-MS analysis shows that knockdown of these transporters enriches acylcarnitines in fly heads just as blocking endocytosis does, supporting the idea that Lrp and Orct are among the proteins affected by the block in endocytosis. Exactly where the accumulation occurs is not known at this time, but we suggest that it is largely extracellular.

Acylcarnitine accumulation in flies with blocked glial endocytosis or lipid transport could occur as a consequence of the high sleep in these animals. We believe this is unlikely as an increase in acylcarnitines appears to generally occur under conditions of sleep deprivation i.e. conditions that would promote sleep. Thus, carnitine conjugation of long chain fatty acids was reported in cortical metabolites of sleep-deprived mice, while short and medium chain fatty acids were reduced (Hinard, Mikhail et al. 2012). Changes in acylcarnitines were also noted in the peripheral blood of sleep-deprived or sleep-restricted humans (Davies, Ang et al. 2014, Weljie, Meerlo et al. 2015) as well as over a day:night cycle (Dallman etal (Ang, Revell et al. 2012, Dallmann, Viola et al. 2012). We find too that a common feature of short-sleeping fly mutants, which are models for chronic sleep deprivation, is an increase in acylcarnitines, regardless of the mechanism that causes the sleep loss (Bedont, Kolesnik et al. 2021). Thus, acylcarnitines are generally associated with sleepiness. While this does not necessarily mean that they promote sleep, we suggest that acylcarnitines are a key marker of sleep need across species, and could be exploited for this purpose.

## Methods

### Fly Stocks

The initial screen was performed with lab stock drivers: ;; Repo-GAL4/TM6c, Sb and UAS-*Dicer*; *Repo*GeneSwitch. SPG driver *Moody-*GAL4 and surface driver 9-137-Gal4 was shared by Roland Bainton, while PG driver NP6293-GAL4 was a gift of Marc Freeman. ;;UAS*-20xShi.ts1* (referred to as UAS*-20xShi^1^*) was shared by Gerald Rubin. *Rab9*-GAL4 (#51587) was acquired from Bloomington. RNAi lines were ordered from VDRC (KK and GD collection) and Bloomington (TRiP collection) stock centers, with the stock number provided in Figure 1 and supplement Figure 1. For control genotypes, GAL4 and UAS lines were crossed to iso31.

### Behavior

Flies were crossed and raised on standard food in bottles. Offspring were kept at 25 ℃, in LD12:12 conditions until at least 6 days post-eclosion, before age-matched flies which were group housed in bottles were used in sleep assays. Mated females were loaded into glass locomotor tubes with 2% agar 5% sugar. Sleep was quantified by the Drosophila Activity Monitor (DAM) system, by the established minimum definition of 5 minutes of inactivity. Data was analyzed in PySolo (Gilestro et al., 2009).

### Metabolomics

Entire flies were quickly frozen in Falcon tubes chilled on dry ice, and placed at −80°C. Each tube contained 50 flies. Heads were subsequently removed from the body by briefly vortexing the tube. Heads were then separated from the rest of the body by an array of copper sieves, whose housing was buried in dry ice to keep the preparation cool. For each sample, 200 fly heads, of equal parts from males and females, were collected in 1.5 mL tubes which were quickly refrozen. Samples were shipped on dry ice to Metabolon, Inc., where they were assessed by LC-MS (Evans...Milgram 2009). For metabolomic analysis of *Lrp* and *Orct* knockdown fly lines, each sample contained 300 fly heads, of equal parts from males and females. Samples were processed by LC-MS at the Penn Metabolomics Core.

### Statistics

For both behavioral and acylcarnitine metabolomics results, the experimental group was compared to two parental controls by One-Way ANOVA with Holm-Sidak post-hoc tests. For the initial comparisons of metabolomics data, Metabolon performed Welch’s t-tests on scaled signal data for each metabolite, between all conditions. Raw signal was scaled so that the median would be equal to 1, using all samples that had been concurrently run. Missing values were filled in with the lowest value of run samples for that metabolite. Additional details of statistics tests are listed in the figure legends.

## Acknowledgement

We thank Dr. Chris Petucci and the Penn Metabolomics Core for providing measurements of acylcarnitines in fly heads.

---

**Table S1:**
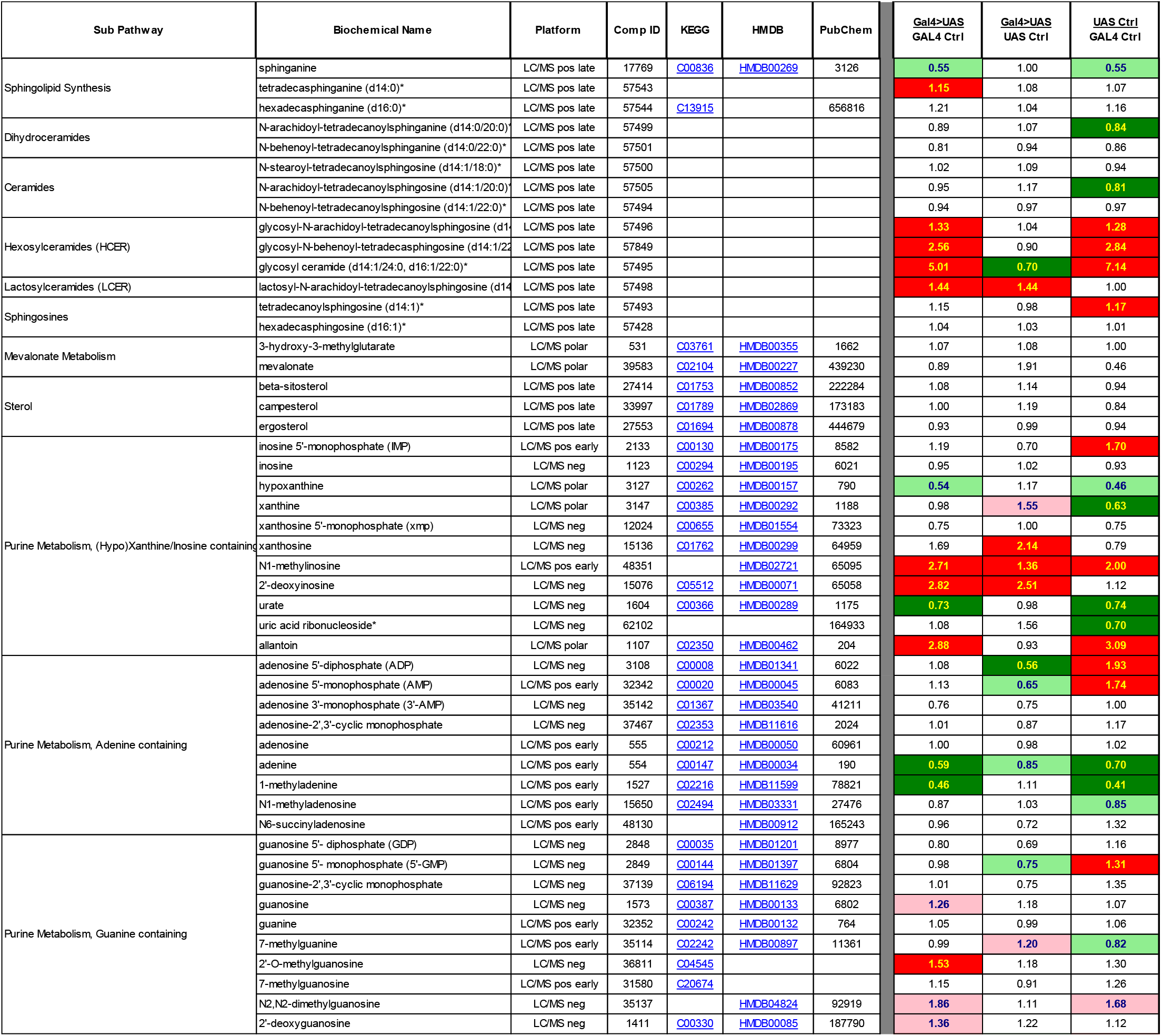
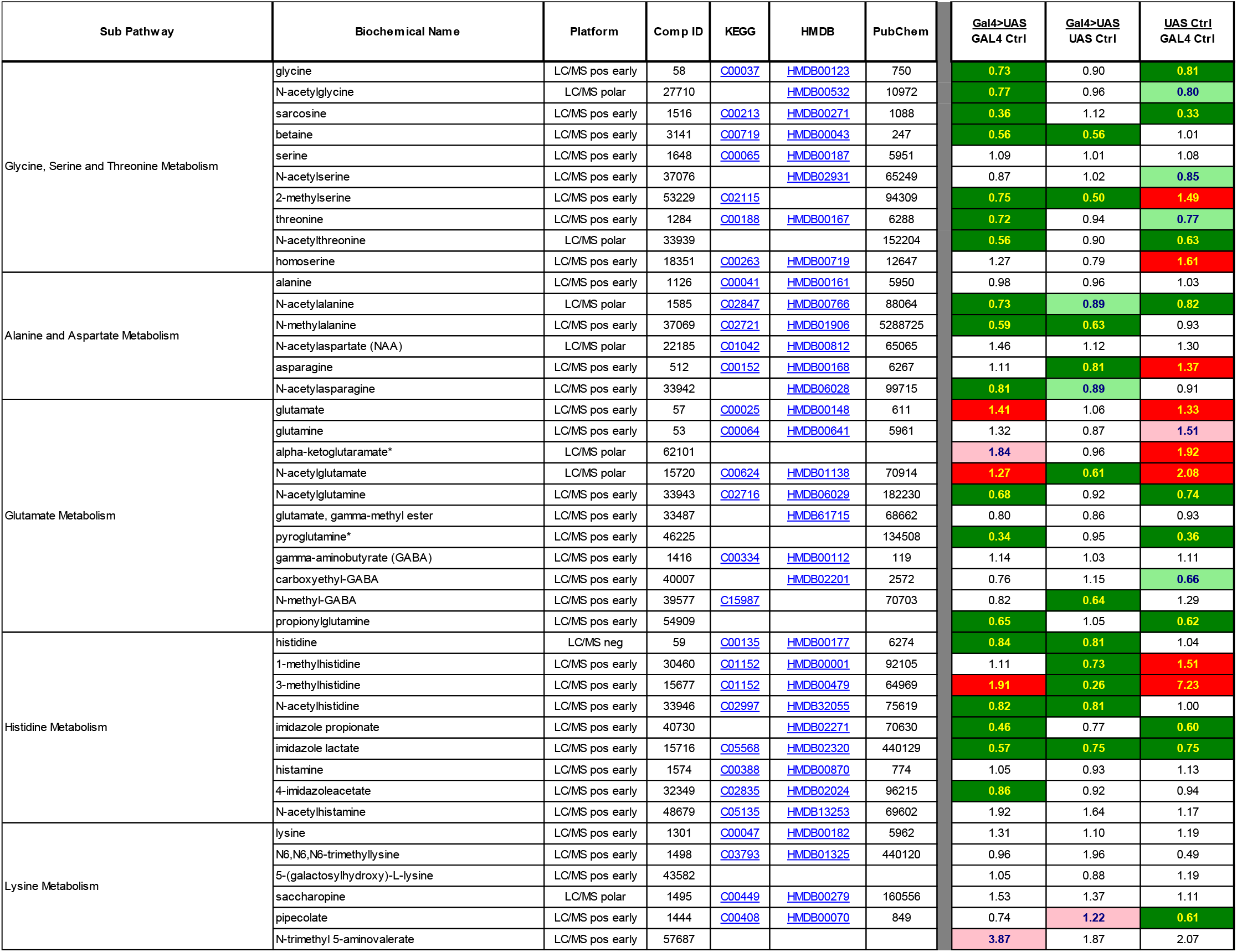
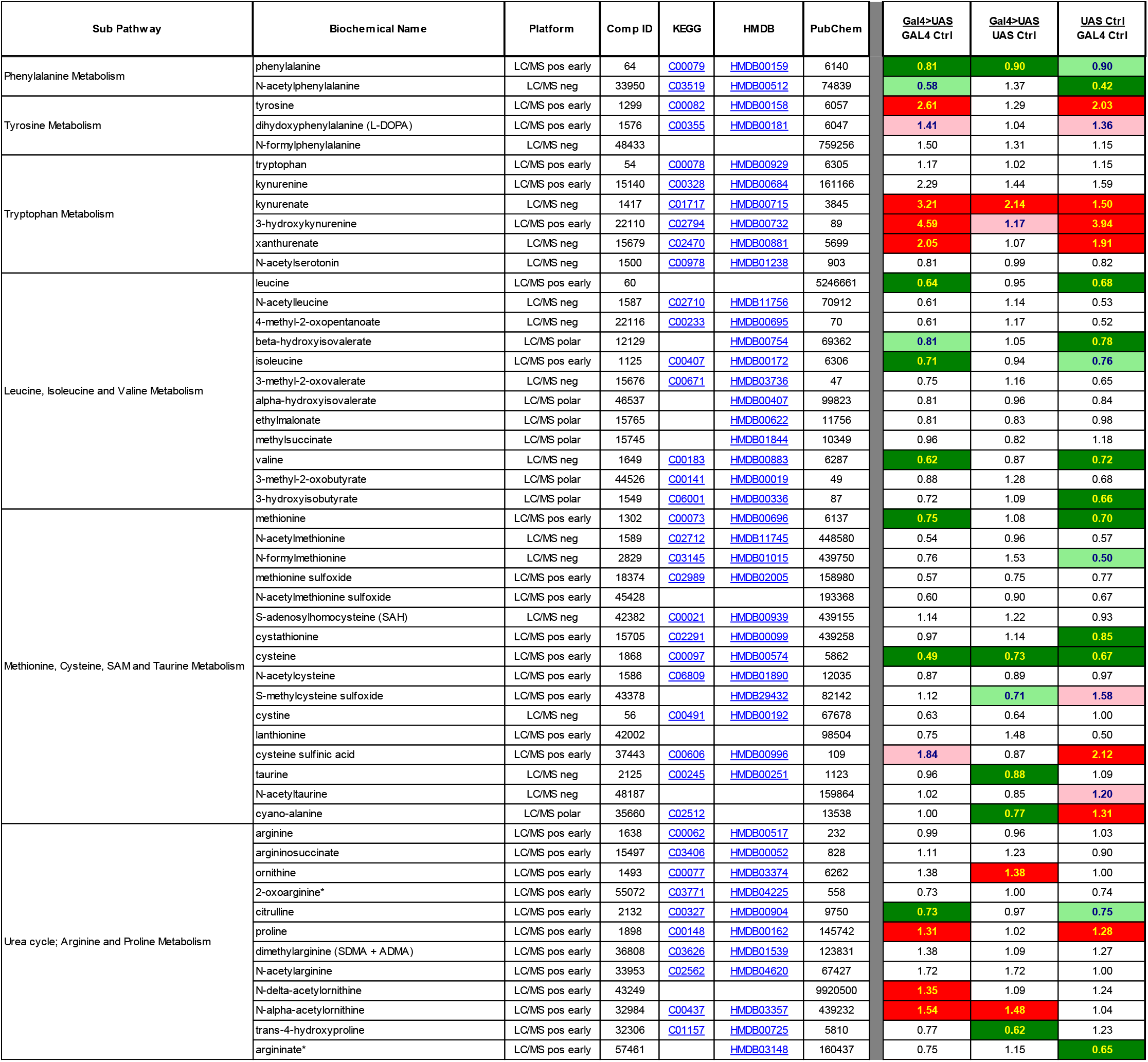
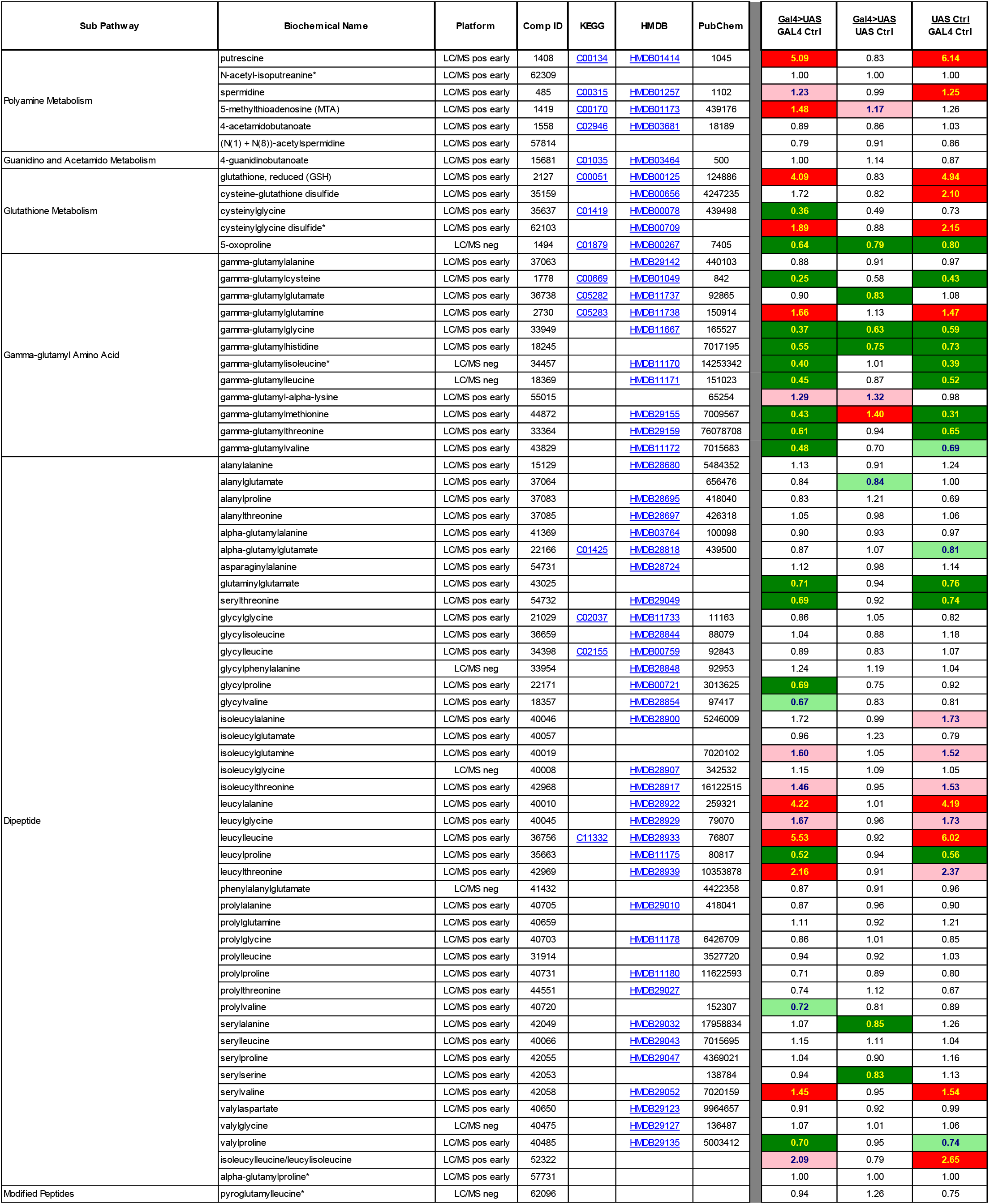
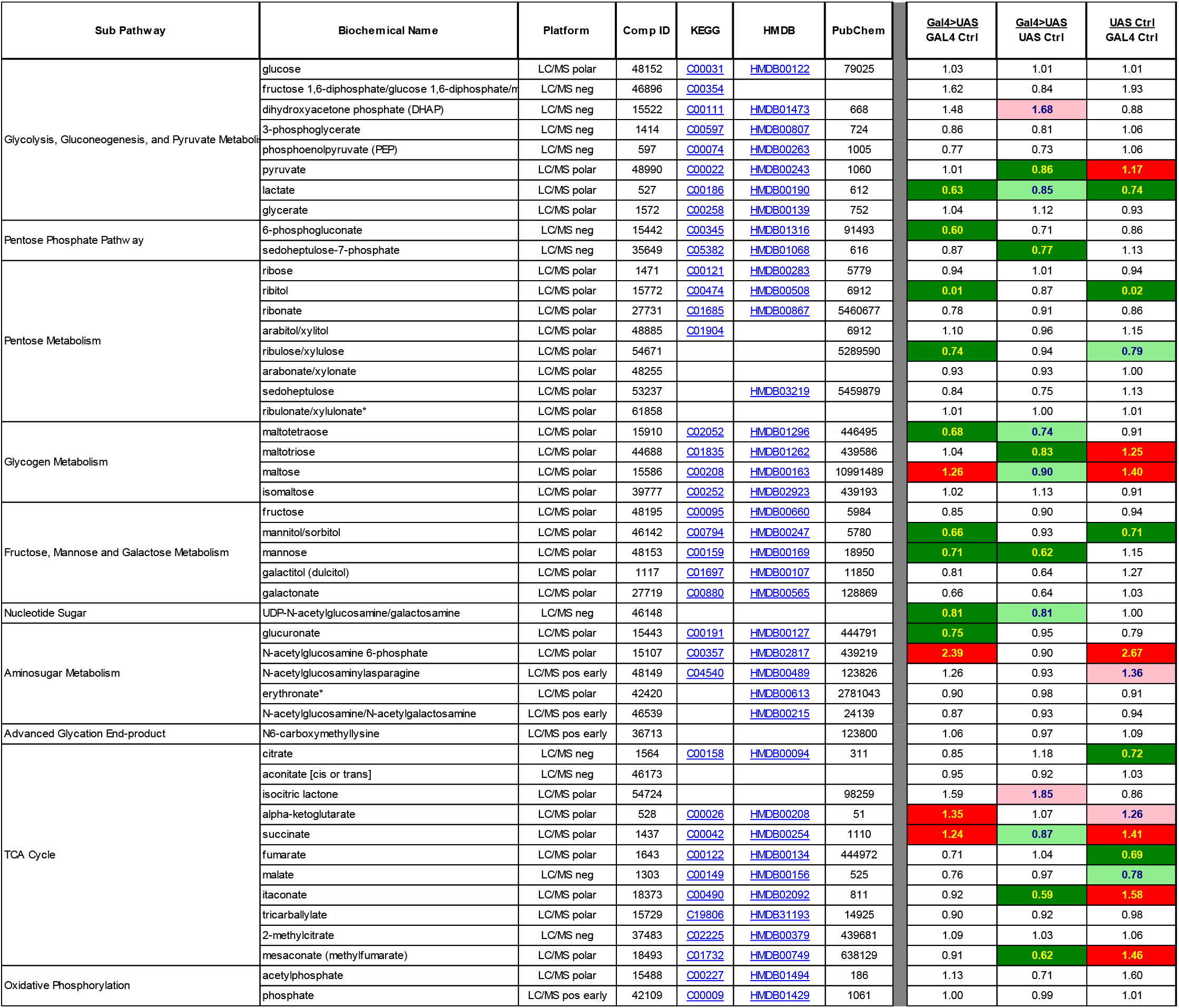
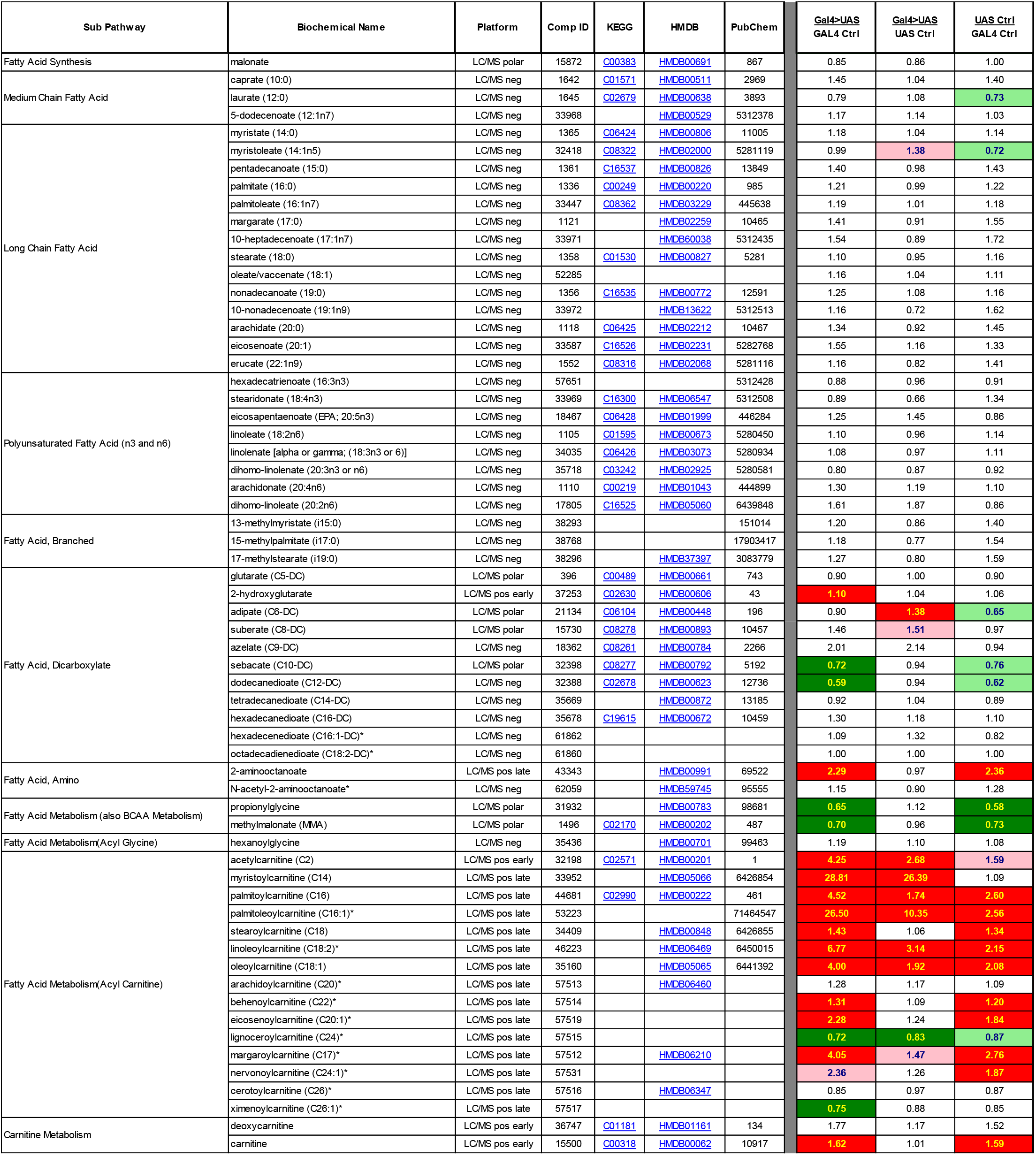
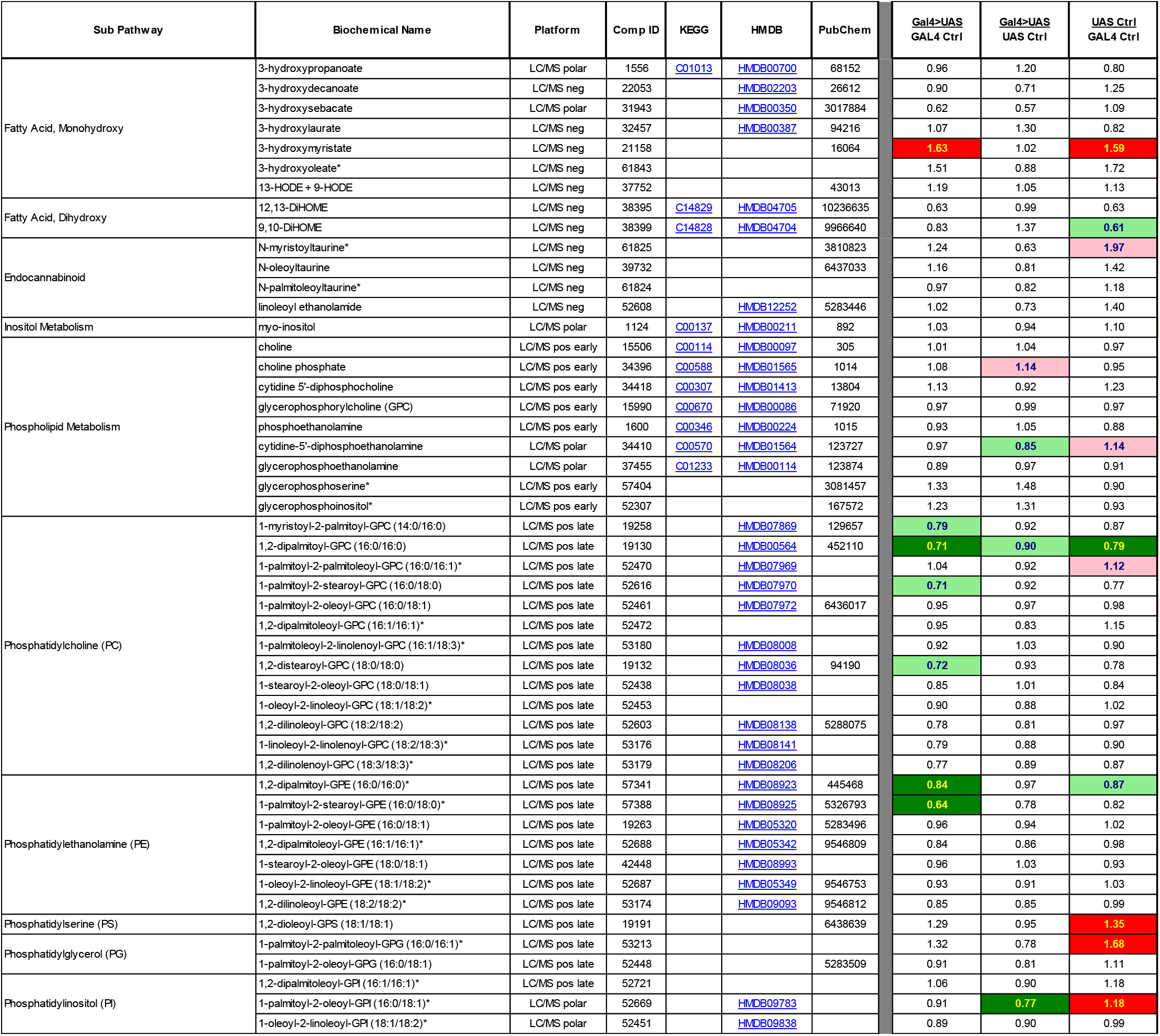
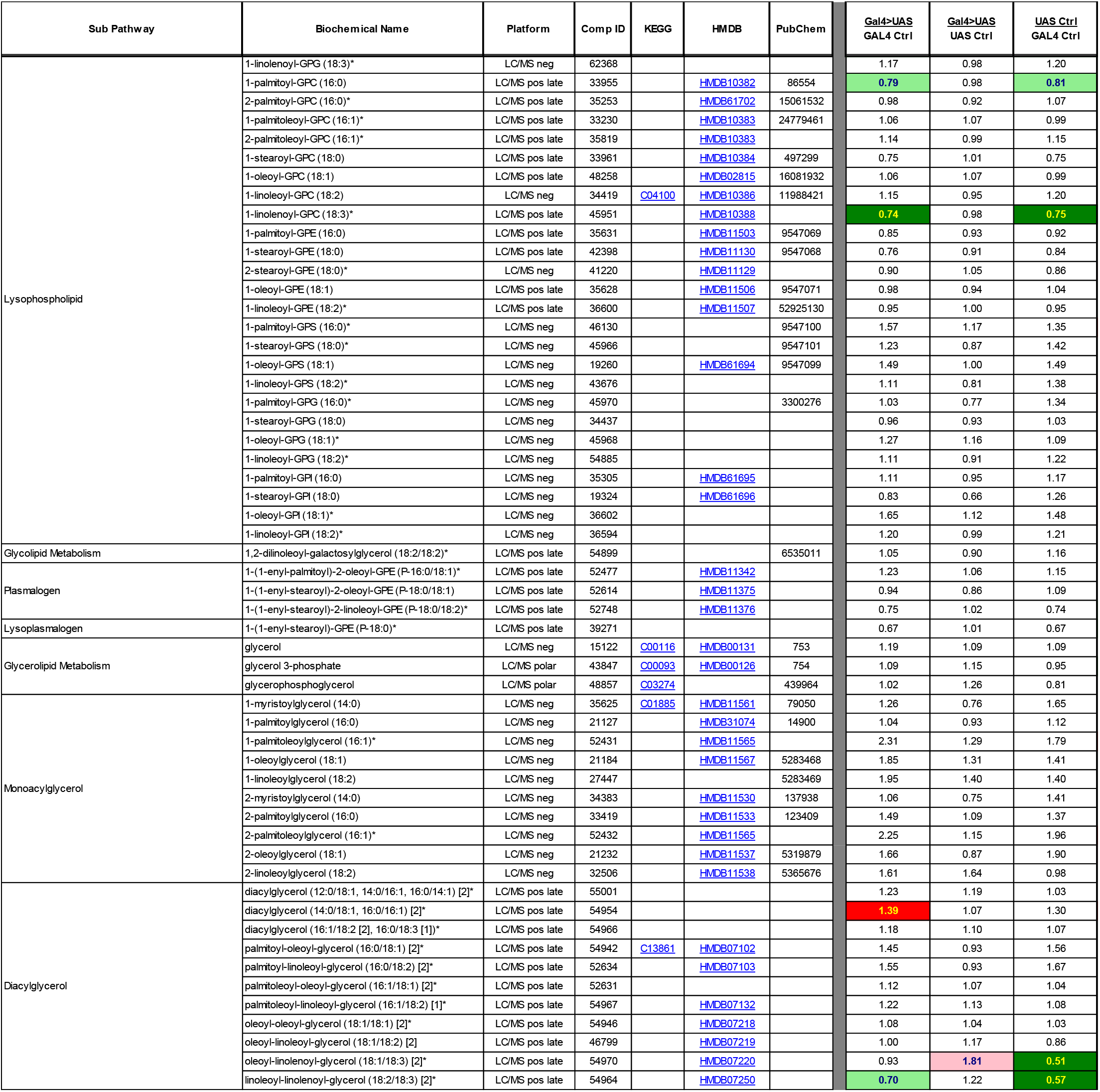
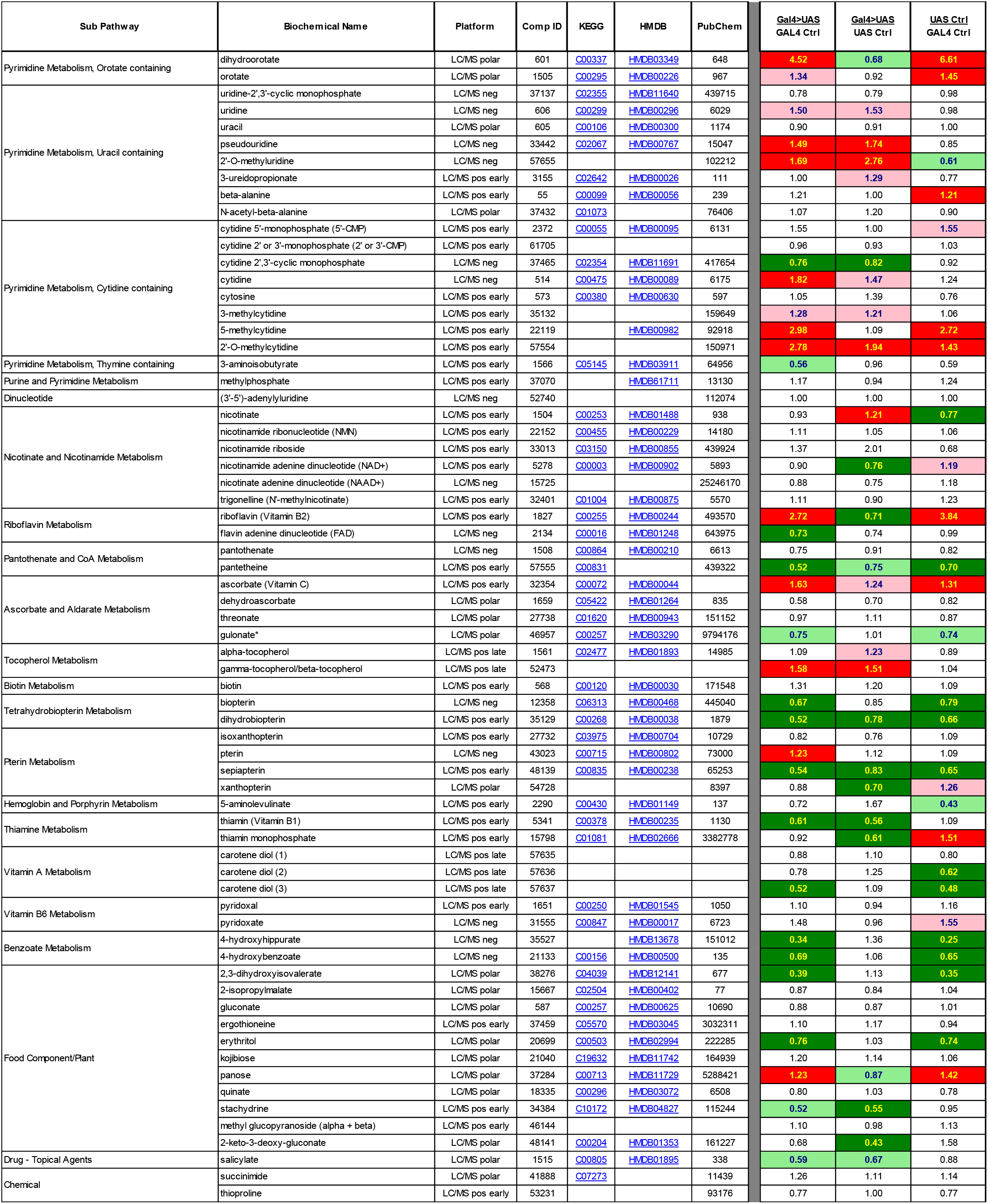
Metabolomics of *Repo*>20x *Shibire* fly heads. All measured metabolites and their respective categories are listed for samples from *Repo*-GAL4>UAS-20x*Shibire*, and both parental controls. Welch’s t-test was performed on scaled signal for each metabolite, comparing each condition. Green highlighting marks a significant difference (p≤0.05) between the groups, where metabolite ratio is < 1.00, while light green is not significant, but close to the threshold (0.05<p<0.10). Red highlighting marks a significant difference (p≤0.05) between groups where metabolite ratio is ≥ 1.00, and light red is not significant, but close to the threshold (0.05<p<0.10).

**Figure 2 supplement 1:**
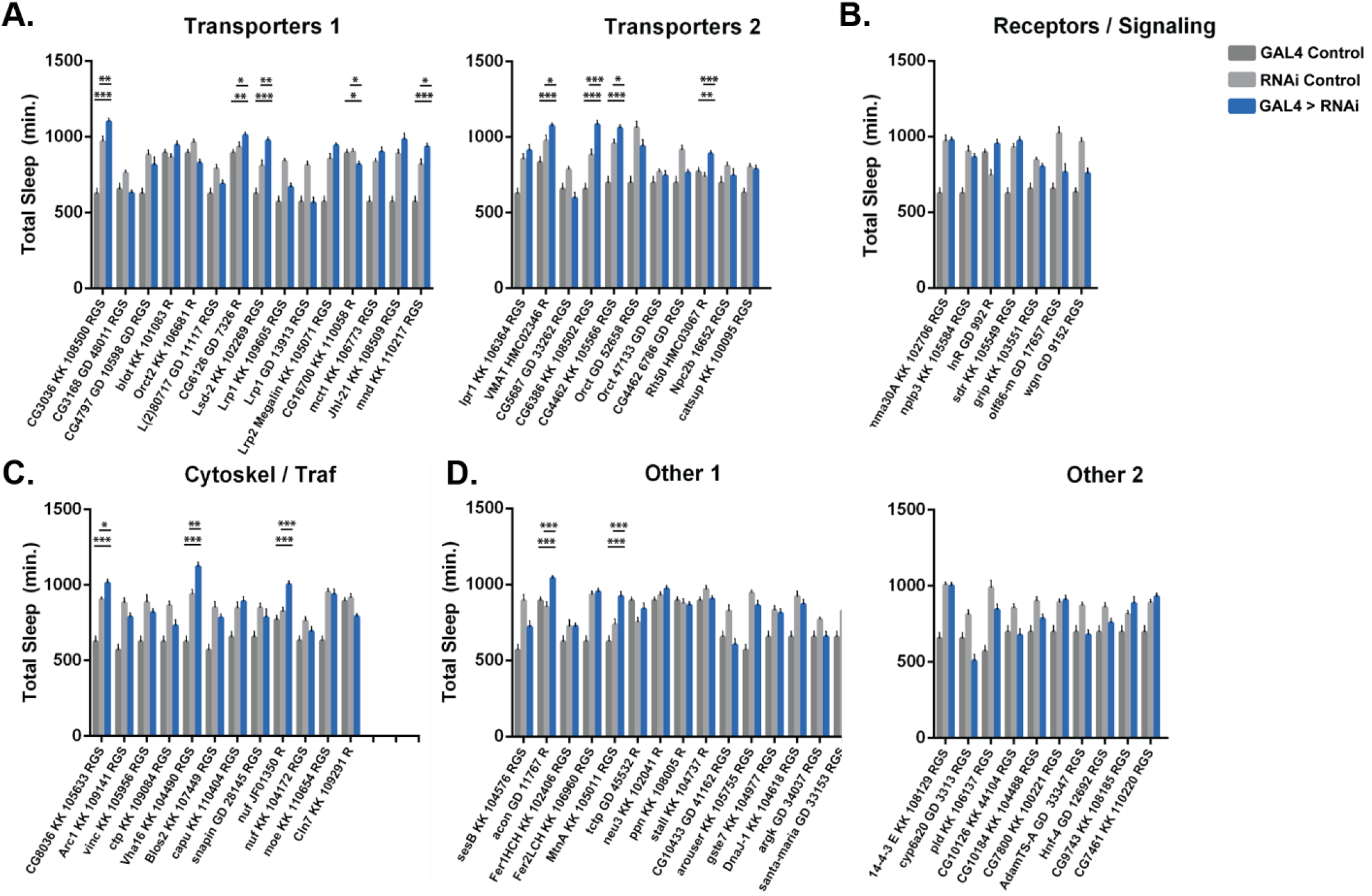
Pan-glial RNAi knockdown screen of candidate genes enriched in the barrier glia. Total sleep in female flies with RNAi knockdown of listed genes (KK and GD are VDRC collections, TRiP lines are from Bloomington Stock Center) (**A**) Transporters, (**B**) Receptors/Signaling pathway factors, (**C**) Cytoskeleton/Trafficking factors and (**D**) other genes by either Repo-Gal4 (labeled R) or UAS-Dicer; RepoGeneSwitch on RU+ food (labeled RGS). n = 9 – 16 flies per genotype, median = 16. One-way ANOVA, with Holk-Sidak post-hoc comparisons. *p < 0.05, **p < 0.01, ***p < 0.001. Error bars represent standard error of the mean (SEM). Significance values only marked for genes in which experimental flies were different from both parental controls. Certain experiments were performed simultaneously and therefore share a Gal4 control, re-plotted per each gene.

**Figure 3 supplement 1:**
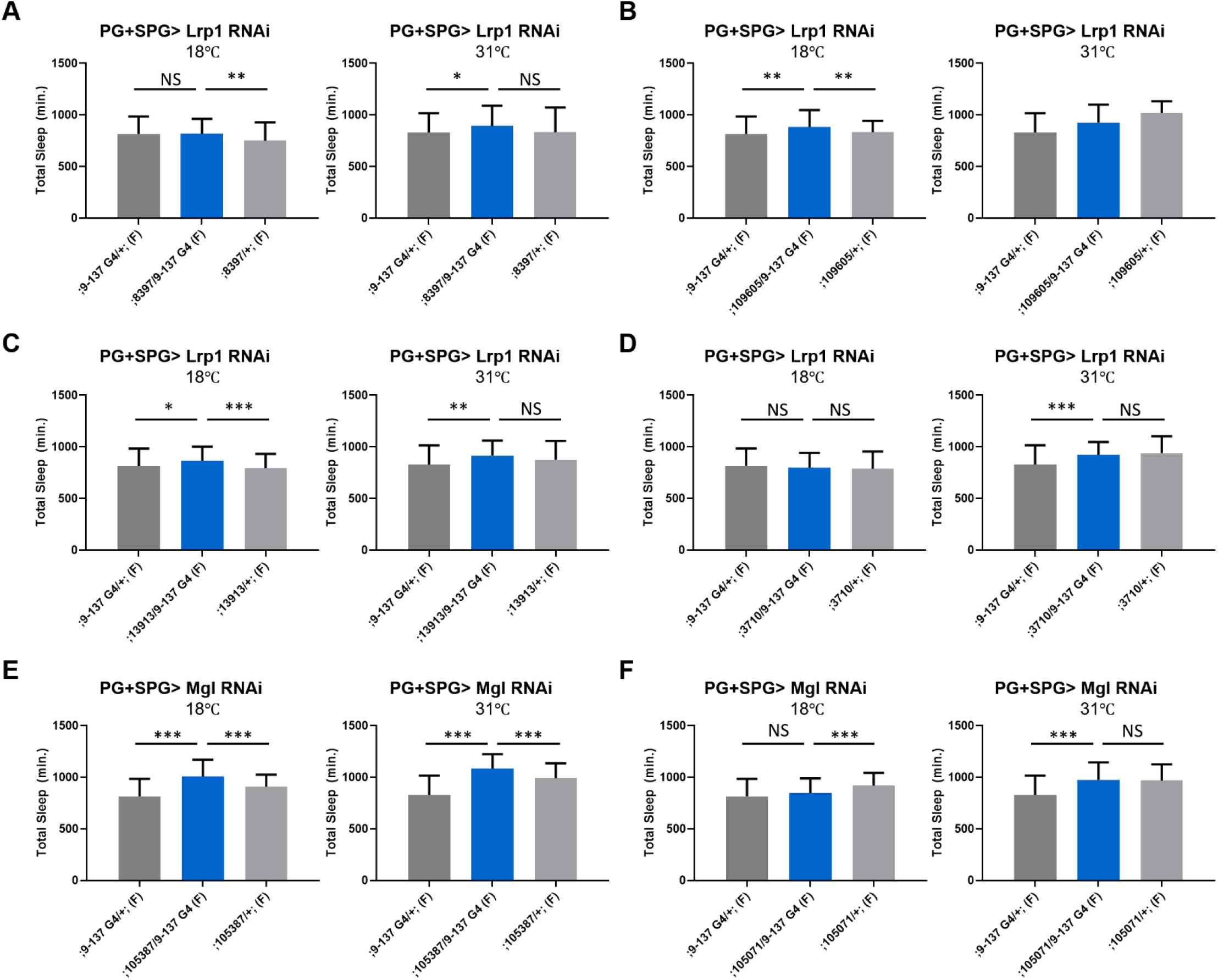
Sleep time changes with knockdown of Lrp genes in surface glia. Total sleep in female flies with knockdown of (**A**) Lrp1 (8397 GD) **(B)** Lrp1(109605 KK) **(C)** Lrp1 (13913 KK) **(D)** Lrp1 (3710 GD) (**E**) Megalin (105387 KK) (F) Megalin (105071 KK) driven by (PG+SPG) driver 9-137-GAL4, n = 40-48 per genotype at 18 ℃(permissive) and at 31 ℃ (restrictive). One-way ANOVA, with Holk-Sidak post-hoc comparisons. *p < 0.05, **p < 0.01, ***p < 0.001. Error bars represent standard error of the mean (SEM).

